# Life on the edge: diet preferences reflect adaptation to drought in *Neotoma fuscipes*

**DOI:** 10.1101/048363

**Authors:** Brendan J Barrett, Arielle Crews, Mary Brooke McElreath

**Author notes:** Corresponding Author: Brendan J Barrett Animal Behavior Graduate Group Department of Anthropology 330 Young Hall One Shields Avenue University of California, Davis Davis, CA 95616.

## Abstract

Ecological change due to habitat fragmentation and climate change can decrease population viability, especially in herbivores and the plant communities upon which they depend. Behavioral flexibility is one important adaptation to both patchy or edge habitats undergoing rapid environmental change. This is true in many generalist herbivores, whose diet preferences can vary substantially, both geographically and over time. Little is known about what plants allow generalist herbivores to respond to rapid environmental changes, and whether these responses are due to variability in diet preference in a population, or individual dietary flexibility. We investigated how the diet preferences of dusky-footed woodrats (*Neotoma fuscipes*) might allow them to respond to drought in a spatially heterogeneous environment. We conducted cafeteria trials on woodrats during a year of extreme drought to assess individual preferences for locally available plants compared to more drought-tolerant edge vegetation. Our results show that woodrats sample a number of plants, but tend to prefer scrub oak, a dominant plant species in the available habitat, as well as chamise- a highly drought-tolerant plant predominantly present in the surrounding edge habitat. No difference in food preferences was detected between sexes, but we found evidence for an effect based on age and proximity to edge habitat. Juveniles who lived closer to the habitat edge were more likely to consume, and consumed greater amounts of plants in cafeteria trials compared to adults and juveniles living further from the edge. In addition to oak and chamise, adults sampled large quantities of other plants such as poison oak and California buckeye, and in general included a wider array of plants in their preferred diets compared to juveniles. We conclude with a discussion of the management implications and outlook for woodrats in the Coast Range of northern California.

---

Climate change may increase the frequency of patchy and edge habitats and alter ecological communities, forcing many species to face rapidly changing, or novel environmental conditions. Rising temperatures, changes in precipitation, and changing hydrology are increasing the frequency, intensity, and duration of droughts in Mediterranean climates worldwide (Cayan et al., 2008; Giorgi and Lionello, 2008). Since 2012, California has experienced exceptional drought conditions, causing declines in many critical sources of water and food. Additionally many plant community ranges are projected to shift in response to climate change (Barbour and Kueppers, 2012), potentially impacting the organisms that depend on them for forage. In response to changing and decreased food availability, one of the primary causes of population declines and extinctions (Cahill et al. 2013), wildlife may either locally adapt or move in search of better habitat. While there is evidence that mammals at higher elevations in California are responding to changing temperatures by shifting ranges, less is known about how populations are responding to climate change in heterogeneous habitats at lower altitudes (Rowe, et al., 2015).

Rather than undergoing range-shifts, species with sufficient behavioral and physiological plasticity may be able to withstand some of the effects of climate change by adding novel foods to the diet (Tuomaninen and Candolin 2011; Varner and Dearing 2013). However, success will depend on how the intensity and duration of change interacts with other variables such as species generation time, recruitment, and predation pressure. From a wildlife management perspective, understanding the extent that behavioral plasticity can permit wildlife to cope with environmental change will help fine-tune and focus conservation efforts. Yet, this kind of information is rarely available to wildlife and land managers. To fill this gap in knowledge and better understand how behavioral plasticity can alleviate the negative effects of climate change-- namely altered food resources --we conducted a study of diet preferences and responses to novel plants in a mammalian population experiencing severe drought conditions.

We focused on dusky-footed woodrats (*Neotoma fuscipes*), a behaviorally plastic herbivore that successfully makes a living in a variety of habitats throughout California and southern Oregon. The species has been identified as an ecosystem engineer (Whitford and Steinberger 2010), due to its construction of large tent-shaped dens that create buffered microclimates not only for woodrats, but also for a long list of species including many arthropods, lizards, snakes, and other small mammals. They are also prey to a number of predators including California spotted owls, coyotes, bobcats, fox, raccoons, skunks, weasels, and snakes. Given their ubiquity and central role in the food web, woodrats are an ideal indicator species (Caro 2010) for California’s Mediterranean ecosystems. Although woodrats are not endangered and are currently classified as “least concern” by IUCN, population declines have been occurring in the species since the early 2000s (McEachern et al. 2007), which may be linked to climate change and increasing drought conditions in California. These factors suggest that woodrats are an excellent system to study how generalist herbivores behaviorally cope with climate change and its effect on food resources.

In particular we were interested in investigating the extent to which edge habitat might provide alternative food sources for woodrats during drought. Edge effects have long been discussed in the wildlife conservation literature, with both positive (e.g., increased biodiversity and abundance) and negative (e.g., increased predation pressure) consequences (Leopold 1933, Lidicker 1999, Kremsater and Bunnell 1999). Previous work on woodrats in Douglas fir / tanoak forests has highlighted this tradeoff; despite increased risk of predation from spotted owls and other predators, woodrats readily cross ecotonal boundaries and frequently use edge habitat (Sakai and Noon 1993, Sakai and Noon 1997). Reasons for this behavior are not well understood, but researchers have speculated that density-dependent factors as well as declines in food quantity or quality are probably involved (Sakai and Noon 1997). Taking this work into account, we included edge effects in our analysis, reasoning that edge habitat may provide alternative food sources for woodrats and affect food preferences during drought. Based on previous diet studies of *N. fuscipes* [(Atsatt and Ingram 1983, McEachern et al. 2006 and the ecologically similar *Neotoma macrotis* (Linsdale and Tevis 1951), formerly a subspecies of *N. fuscipes* (Matocq 2002)], we expected to find that woodrats prefer locally available plants, such as scrub oak. However, we also hypothesized that proximity to edge habitat mediates this effect—individuals living close to edge habitat may be more likely to sample edge plants, and novel plants in general—simply because they have more experience with plant variety. Our results quantify behavioral plasticity in food preferences in woodrats along an ecotonal gradient, and examine the value of edge habitat to wildlife conservation, particularly during times of environmental stress. As our results suggest, proximity to edge habitat may enhance woodrat survival during drought, when access to alternative food resources can be extremely beneficial.

## MATERIALS AND METHODS

### Study Site

Experimental trials were conducted June- September 2013 at the Quail Ridge Reserve, in Napa County, California (38°49’ N, 122°14’W), part of the UC Davis Natural Reserve System. 2013 was a significant drought year, the driest year in California’s recorded history, and the second year of drought in the state’s current dry period, in its fifth year as of 2016. A population (N=22) of dusky-footed woodrats (*Neotoma fuscipes*) living in mixed oak woodland habitat bordered by chamise chaparral was live-trapped for inclusion in the study. 9 of these 22 individuals were run multiple times through cafeteria trials, in order to examine the extent of variation that may occur within the same individual. The study site (area=0.93 hectares) was located on a north-facing slope, dominated by California scrub oak (*Quercus berberidifolia*), poison oak (*Toxicodendron diversilobum*), toyon (*Heteromeles arbutifolia*), California bay laurel (*Umbellularia californica*), and California buckeye (*Aesculus californica*). Chamise (*Adenostoma fasciculatum*) dominated the edge and surrounding habitat on south facing slopes. The two habitat types were distinctly separated by an approximately 2 m wide fire road along the spine of the ridge (hereby referred to as the habitat edge).

### Live-Trapping

The study area was regularly live-trapped 1 night per month to mark and monitor the woodrat population. Trapping effort included 2 XLK Sherman traps (7.62 x 9.525 x 30.48 cm; H.B. Sherman Traps, Inc) baited with oats and placed directly outside each woodrat den. Traps were set at dusk and checked the next morning shortly after sunrise. All woodrats were sexed, weighed, and marked with Biomark HPT12 pit tags, as well as standard metal ear tags. For cafeteria trials, traps were set an additional 2 - 3 nights per month. Woodrats included in cafeteria trials were transferred to separate holding cages and transported a short distance (< 500 m) to an outdoor field “laboratory” (large tent equipped with a table, experimental arena, scale, and video camera). During the day, woodrats were held under the shade of a large tent, and their holding cages were covered with a cotton sheet to simulate the dark environment of their dens. They were provided with water ad libitum and deprived of food until nightfall to encourage participation in cafeteria trials.

### Cafeteria Trials

Cafeteria trials were conducted at night (20:00-01:00) when woodrats are typically active and foraging. In order to examine the dietary preferences of *N. fuscipes*, trapped individuals were presented with a total of 7 plant foods. These included 5 plants common in the immediate habitat and familiar to all woodrats in our study: bay, oak, toyon, buckeye, and poison oak. In addition we included chamise, a drought-tolerant plant that dominates the surrounding edge habitat. Chamise was potentially novel to woodrats in our study living in interior oak woodland habitat. Thus, to establish a baseline to responses to novel food, as well as investigate any patterns in that response (e.g, by age or sex), we also included 1 completely novel plant sample, incense cedar (*Calocedrus decurrens*). Incense cedar is naturally consumed by other woodrat populations in California (McEachern et al. 2006), but is not present at the Quail Ridge Reserve. Like chamise and all of the plants included in our study, incense cedar is chemically defended with plant secondary compounds.

Fresh plant samples were offered simultaneously and in equal amounts (2 g- wet weight). Trials were run individually, in glass experimental arenas (0.6 x 0.3 x 0.3 m) with wire mesh lids. For each trial, a single woodrat was introduced into the arena for an initial acclimation period of 10 minutes. After acclimation, a glass partition was used to confine the woodrat while the seven plant samples were placed in the arena in randomized order. The woodrat was then allowed to sample plants for 15 minutes under video surveillance. Following trials, the remaining uneaten plant samples were removed from the arena, weighed, and recorded. All woodrats were released at their points of capture (i.e. den) following cafeteria trials. All animal procedures followed ASM guidelines (Sikes et al. 2011) and were conducted under UC Davis IACUC approval (Protocol 16253).

### Distance from Edge Habitat

The UTM location and altitude of woodrat dens were marked using a handheld GPS unit (Garmin GPS MAP 64s). A fire road marking the habitat boundary between the north-facing oak forest slope form the south-facing chamise slope was mapped using GPS points every 1 m. Distance between a den and habitat boundary was calculated as the minimum two-dimensional distance between the two points. Analyses were also run with three-dimensional Euclidean distance and yielded similar results, however these were not used in the final analysis due to large measurement errors in elevation.

### Statistical Analysis

Data were analyzed using a series of generalized linear mixed models in R (v. 3.2.1) using the map2stan function in the rethinking package (McElreath, 2015) and Hamiltonian Monte Carlo implementation in RStan v. 2.70 (Stan Development Team, 2015). As distances are positive continuous variables, which are well described by a gamma distribution, we used a gamma-distributed GLM to model the distances of adult and juvenile woodrat dens relative to the habitat edge. For the diet preference models, 6 models were fit to the observed data and compared using Widely Applicable Information Criterion (WAIC) (Table 2). WAIC is a generalized form of information theoretic model comparison (of which the more familiar AIC is a special case), suited for Bayesian hierarchical model comparison (Watanabe 2010). All models for the cafeteria trial study assumed a zero-augmented gamma distribution for the outcome variable (grams of plant material eaten). A zero-augmented gamma is a mixture distribution that combines a Bernoulli and gamma distribution. The Bernoulli component estimates *p*, the probability of an individual consuming a plant (measured as a 1/0 outcome). The gamma component estimates *μ*, the amount of food eaten given that an animal chooses to sample food. Model estimates for *p* and *μ* were used as metrics for diet preference. Predictor variables included age, sex, distance to the habitat edge (log transformed), and an interaction between age and distance. We included varying slopes for each plant species consumed for these predictors, and varying intercepts for individual woodrats, plant species, experimental date, and experimental trial for each individual (if they were assayed multiple times). Using varying effects statistically accounts for concerns of psuedoreplication (Schank and Koehnle 2009) and unequal sampling across experimental dates and individuals. Instead of p-values, we present posterior mean estimates (PME) of the parameters *p* and *μ* (converted to probabilities and grams respectively), posterior distribution samples, and 90% posterior credible intervals (CI) to communicate effect size, variation, and uncertainty.

## RESULTS

WAIC model weight provides a means to assess the value of different combinations of predictors in out-of-sample prediction. We considered models with main effects including distance from edge, sex, age, and age by distance and sex by distance interaction effects. WAIC favored the model (m6, wWAIC = 91%) that included distance, age, and an interaction between age and distance (Table 1). Summaries of the parameters estimated in the models and WAIC values for m6 can be seen in Table 2.

**Table 1.** Estimated parameters and WAIC values of all models fit. Models are listed in order of lowest WAIC value. dWAIC indicates the difference between the WAIC value of listed model and the highest ranked model, while WAIC weight is the proportion of posterior probability assigned to each model. \**All models listed contained the following parameters as varying intercepts: α*_*species*_, *α*_*ratidt*_, *α*_*order*_, *α*_*date*_.

| Model | Fixed Effects | Random Effects* | WAIC | dWAIC | wWAIC |
| --- | --- | --- | --- | --- | --- |
| m6 | $\alpha$, $\beta$ distance, $\beta$ adult, $\beta$ adult*distance | $\beta$ distance (species), $\beta$ adult (species) | -252.25 | 0.00 | 0.91 |
| m5 | $\alpha$, $\beta$ distance, $\beta$ adult | $\beta$ distance (species), $\beta$ adult (species) | -246.99 | 5.25 | 0.07 |
| m4 | $\alpha$, $\beta$ adult | $\beta$ adult (species) | -243.04 | 9.21 | 0.01 |
| m3 | $\alpha$, $\beta$ distance | $\beta$ distance (species) | -242.66 | 9.59 | 0.01 |
| m2 | $\alpha$, $\beta$ distance, $\beta$ adult | $\beta$ adult (species) | -241.79 | 10.46 | 0.00 |
| m1 | $\alpha$, $\beta$ male | $\beta$ male (species) | -238.76 | 13.49 | 0.00 |

**Table 2.** Parameter estimates including posterior means, standard deviations, and 90% credible intervals (CI) estimated from the highest ranked model (m6).

| Parameter | Posterior Mean | Std Dev | 90% CI |
| --- | --- | --- | --- |
| Fixed Effects |  |  |  |
| $\alpha$ | -0.87 | 0.62 | -1.94 , 0.07 |
| $\beta$ distance | -0.20 | 0.14 | -0.42 , 0.04 |
| $\beta$ adult | -1.36 | 0.61 | -2.35 , -0.36 |
| $\beta$ adult*distance | 0.40 | 0.19 | 0.09 , 0.71 |
| scale | 0.12 | 0.01 | 0.1 , 0.14 |
| Random Effects |  |  |  |
| $\alpha$ species | | | |
| $\alpha$ poison oak | -0.26 | 0.48 | -1.05 , 0.51 |
| $\alpha$ toyon | -0.70 | 0.53 | -1.53 , 0.14 |
| $\alpha$ bay laurel | -0.47 | 0.51 | -1.29 , 0.34 |
| $\alpha$ buckeye | -0.30 | 0.48 | -1.08 , 0.48 |
| $\alpha$ incense cedar | -0.56 | 0.50 | -1.39 , 0.21 |
| $\alpha$ chamise | -0.05 | 0.46 | -0.81 , 0.68 |
| $\alpha$ scrub oak | 1.52 | 0.55 | 0.6 , 2.38 |
| Varying Slopes |  |  |  |
| $\beta$ distance (species) | | | |
| $\beta$ distance (poison oak) | 0.05 | 0.13 | -0.16 , 0.26 |
| $\beta$ distance (toyon) | 0.13 | 0.15 | -0.08 , 0.36 |
| $\beta$ distance (bay laurel) | 0.08 | 0.14 | -0.13 , 0.3 |
| $\beta$ distance (buckeye) | 0.04 | 0.13 | -0.17 , 0.24 |
| $\beta$ distance (incense cedar) | 0.05 | 0.13 | -0.15 , 0.28 |
| $\beta$ distance (chamise) | 0.02 | 0.12 | -0.18 , 0.22 |
| $\beta$ distance (scrub oak) | -0.25 | 0.16 | -0.48 , 0.02 |
| $\beta$ adult (species) | | | |
| $\beta$ adult (poison oak) | 0.05 | 0.18 | -0.24 , 0.32 |
| $\beta$ adult (toyon) | 0.04 | 0.18 | -0.22 , 0.36 |
| $\beta$ adult (bay laurel) | 0.10 | 0.18 | -0.15 , 0.41 |
| $\beta$ adult (buckeye) | 0.06 | 0.18 | -0.22 , 0.35 |
| $\beta$ adult (incense cedar) | 0.06 | 0.18 | -0.22 , 0.35 |
| $\beta$ adult (chamise) | 0.02 | 0.17 | -0.24 , 0.3 |
| $\beta$ adult (scrub oak) | -0.24 | 0.22 | -0.6 , 0.06 |
| Random Intercepts Variance |  |  |  |
| $\sigma$ ratid | 0.13 | 0.08 | 0.01 , 0.24 |
| $\sigma$ order | 0.71 | 0.71 | 0.02 , 1.47 |
| $\sigma$ date | 0.38 | 0.21 | 0.1 , 0.67 |
| $\sigma$ [$\alpha$ species] | 0.86 | 0.36 | 0.33 , 1.34 |
| $\sigma$ [$\beta$ distance (species)] | 0.22 | 0.16 | 0.01 , 0.41 |
| $\sigma$ [$\beta$ adult (species)] | 0.17 | 0.10 | 0.01 , 0.29 |

### Overall diet preference

We defined preference based on a mean percent of diet greater than 14.3%, which represents the percent consumed if all plants were consumed in equal quantity. Pooling data over all individuals, woodrats demonstrated a strong preference for scrub oak, which comprised 39 % of the overall diet. Models predicted that woodrats had a higher probability of sampling oak (*p* = 0.30 [90% CI = 0.17 - 0.51]; Fig 1g), and consumed larger quantities of it (*μ* = 0.49 g [90% CI = 0.21 - 1.05]; Fig 2g) compared to alternative plants. The second most abundantly consumed food was chamise in both the probability of being sampled (*p* = 0.19 [90% CI = 0.09 - 0.36]; Fig 1f) and amount consumed (*μ* = 0.25 g [90% CI = 0.10 - 0.55]; Fig 2f). Chamise is only available at the habitat edge, and is the dominant species on the slope opposite our study site, yet it contributed 17% of the overall diet. California buckeye accounted for 14.7% of the diet, indicating a very slight preference. Cedar, the novel food source, was the least sampled food with respect to probability of being sampled (*p* = 0.07 [90% CI = 0.06 - 0.26]; Fig 1e) and the amount consumed (*μ* = 0.16 g [90% CI = 0.07 - 0.36]; Fig 2e), corresponding to 4.1% of the diet.

**Figure 1.**
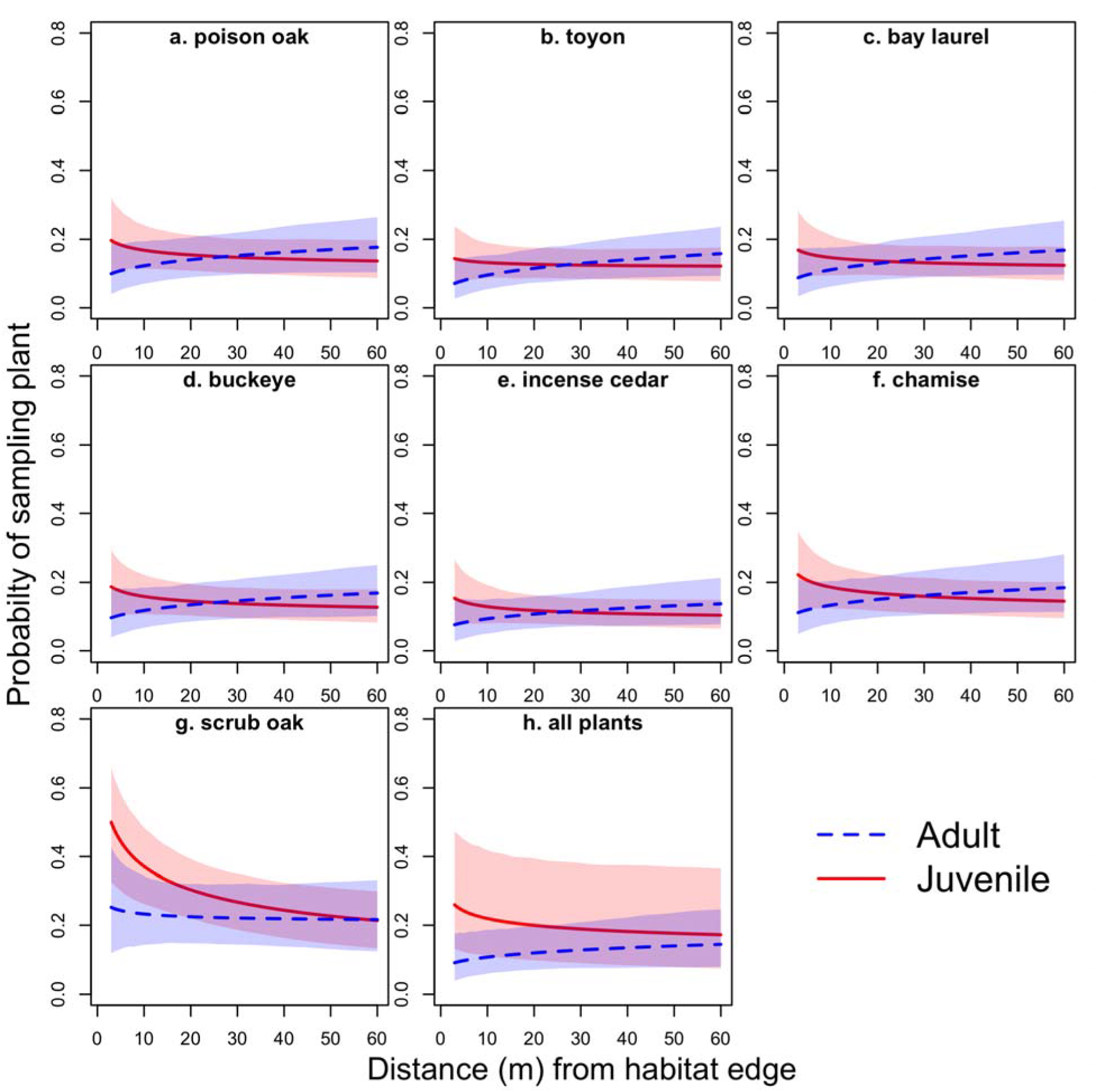
Probability of consuming plant vs. distance from habitat edge. These graphs show model averaged predictions for *p*- the probability that a plant was consumed, for each plant species in the cafeteria trial dependent upon if consumed by an adult (blue dashed lines) or juvenile (red solid lines). Lines are estimates of posterior means presented along 90% confidence intervals. Dark grey regions indicate where CIs overlap.

**Figure 2.**
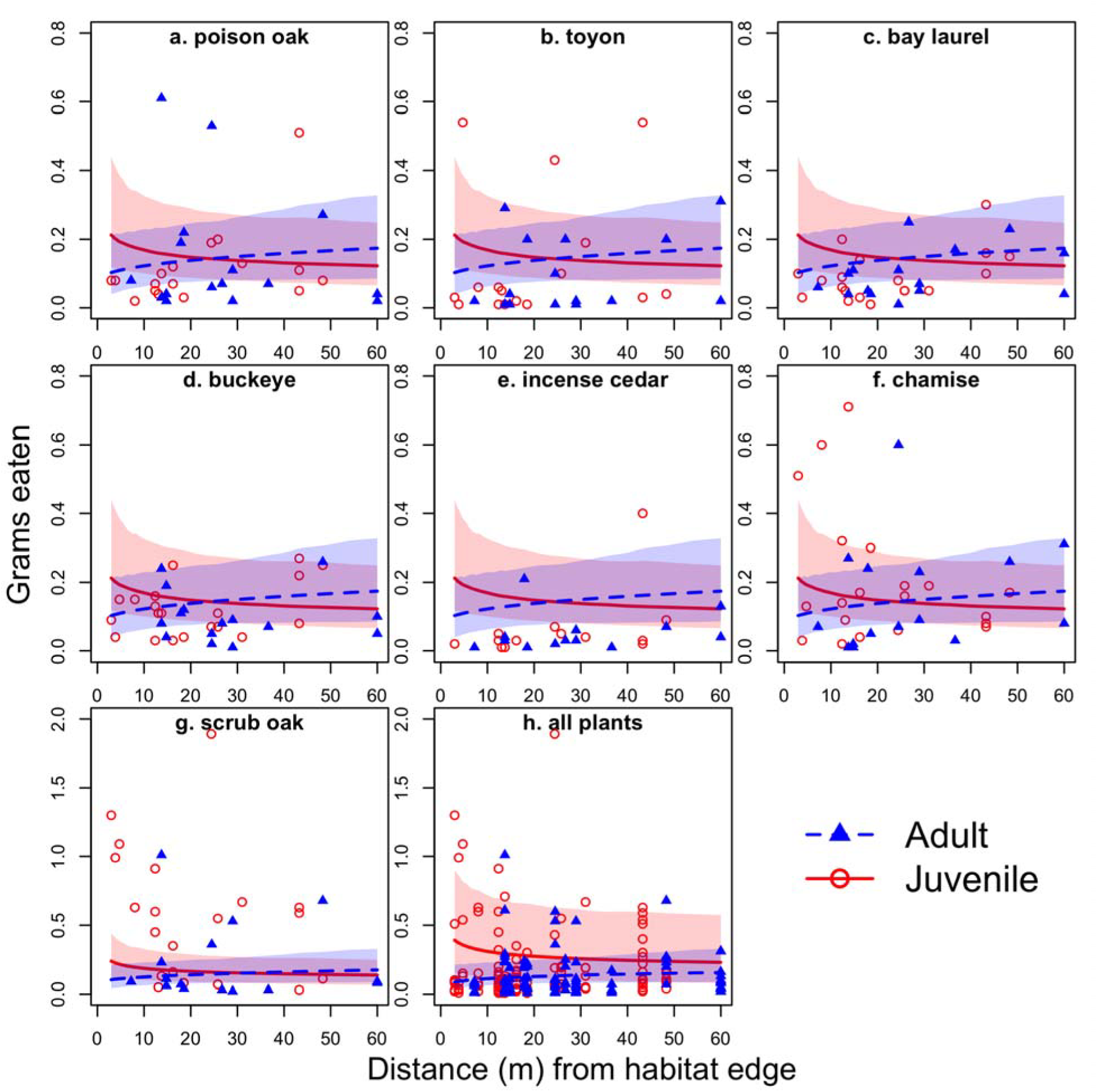
Mass of plant material consumed vs. distance from habitat edge. These graphs show model averaged predictions for α - the mass of plant material consumed for each plant species in the cafeteria trial dependent upon if consumed by an adult (blue dashed lines) or juvenile (red solid lines). Lines are estimates of posterior means presented along 90% confidence intervals. Dark grey regions indicate where CIs overlap. Points are all measures instances of plant consumption > 0 grams.

### Age differences

Overall, juveniles were slightly more likely to consume plant material than adults (*p*_Juvenile_ = 0.20 [90% CI = 0.10 - 0.38]; *p*_Adult_ = 0.17 [90% CI=0.08 - 0.33]; Fig 1h), and consumed a greater amount of plant material across all plant species (β_adult_ = −1.36 [90% CI = −2.35 - −0.36]; *μ*_Juvenile_ = 0.27 g [90% CI = 0.11 - 0.61]; *μ*_Adult_ = 0.22 g [90% CI = 0.09 - 0.49]; Fig 2h). In terms of mean percent of diet, scrub oak accounted for only 25.8% of the diet in adults, whereas it contributed 46.2% in juveniles. Adults tended to sample less scrub oak and included more of the other 6 plant species in their diets compared to juveniles (Table 3 and Figure 2).

**Table 3.**
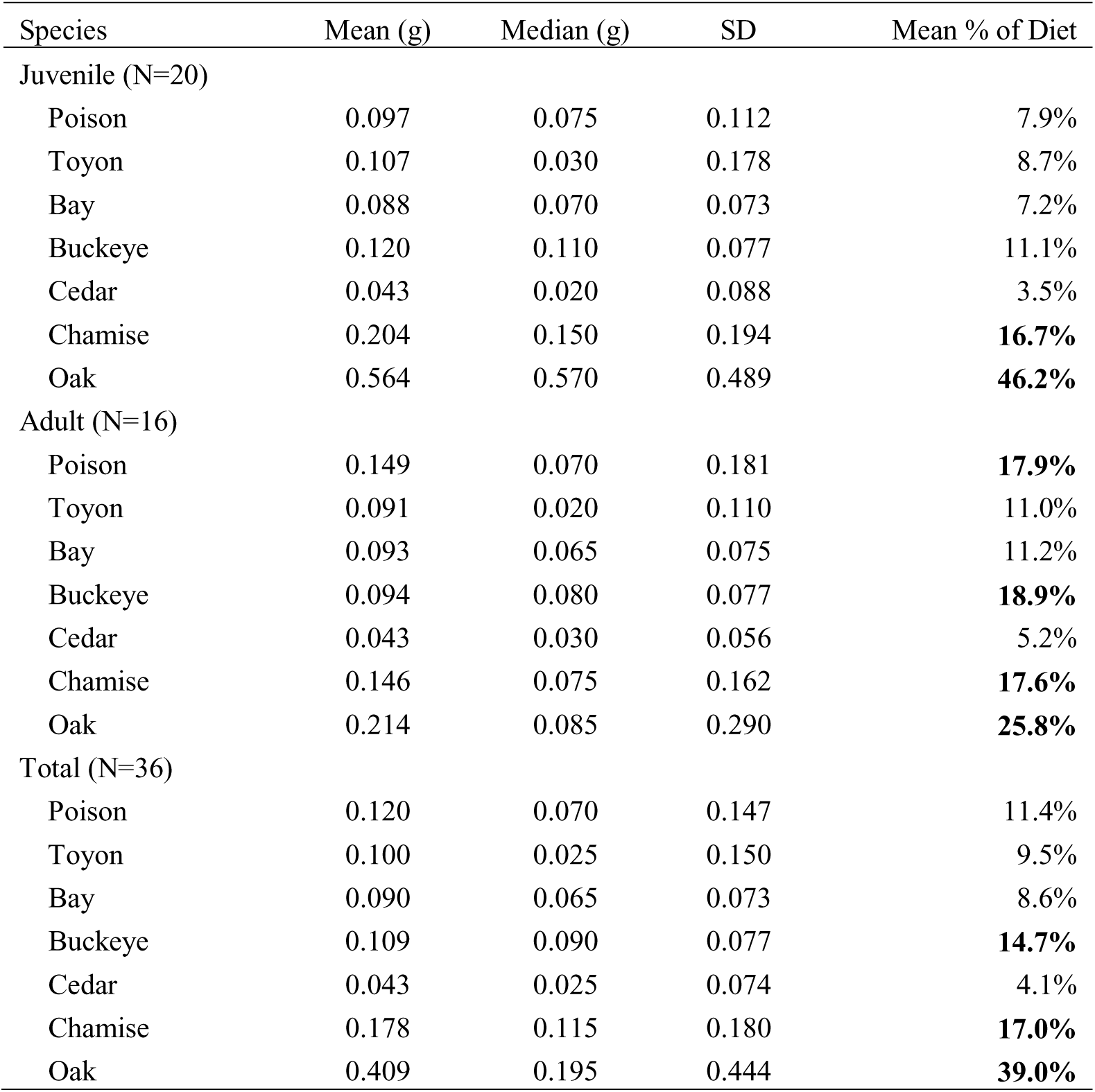
Summary statistics for the average amounts of each plant species sampled.

Estimates from the GLM comparing the distance of woodrat dens relative to the habitat edge suggest that juveniles are more likely to live closer the edge (PME_juvenile_ = 18.62 m [90% CI = 14.99 - 22.01]) than adults (PME_adult_ = 26.33 m [90% CI = 21.43 - 31.21]). However the overlap of posterior distributions (Fig 3) for adults and juveniles does not provide clear support for a solely age-based effect. Instead, dietary preferences for adults and juveniles should be considered in light of their proximity to edge habitat, as discussed in the next section.

**Figure 3.**
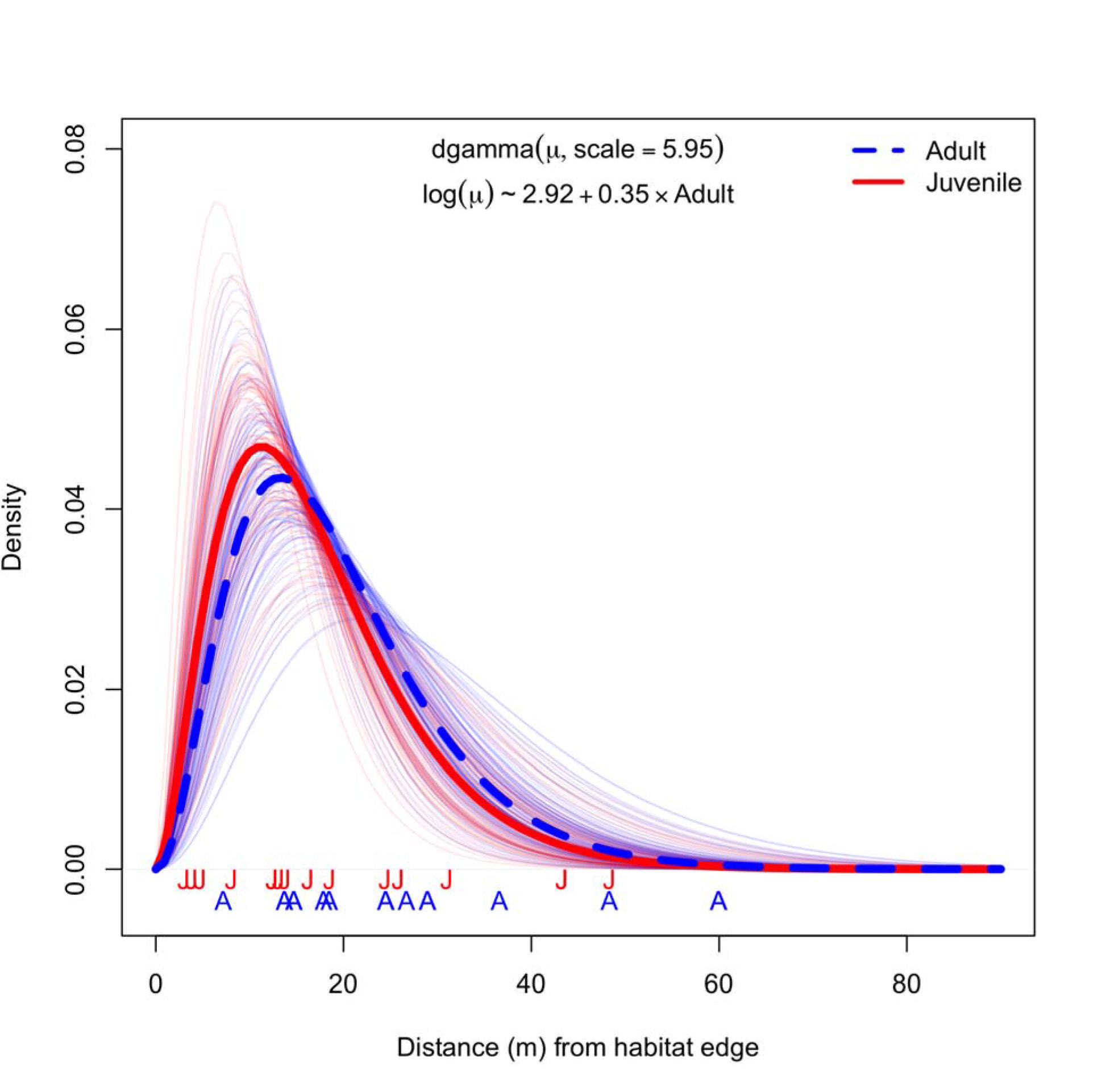
Posterior distributions from a gamma generalized linear model of the abundance of woodrat houses relative to the habitat edge. Thicker lines are posterior medians for adults (blue dashed) and juveniles (red), and thinner lines are 100 posterior samples drawn for adults and juveniles to communicate uncertainty about estimates. The regression equation for this model can be seen at the top of the figure.

### Distance effects

Location of woodrat dens relative to the habitat edge (β_distance_ = −0.20 [90% CI = −0.42 - 0.04]) and the interaction between woodrat age and distance from habitat edge (β_adult*distance_ = 0.40 [90% CI = 0.09 – 0.71]) were the most important predictors in explaining diet preference at our study site, as indicated by WAIC scores and parameter effect sizes. For juveniles across all plant types, a negative relationship between distance from habitat edge and grams eaten (Fig 2a-h) and distance from habitat edge and probability of a plant being consumed were observed (Fig 1a-h). In contrast to juveniles living far from the habitat edge, adult woodrats further from the edge were more likely to consume each plant item and ate more of each food item, with one exception- scrub oak. Adults also showed a near zero relationship between scrub oak consumption and distance to habitat edge, whereas juveniles showed a strong negative relationship (Fig 1g & 2g).

### Sex differences

Male and female woodrats consumed similar amounts of food (PME_male_ = 0.29 g [90% CI = 0.09 - 0.48]; PME_female_ = 0.30 g [90% CI = 0.08 - 0.50]), and displayed similar preferences across plant species. Models including sex were not well ranked according to WAIC values.

### Temporal and order effects

Varying intercept estimates of each trapping session show no noteworthy pattern for how much plant material woodrats consumed. However, woodrats that underwent multiple trials consumed more plant material in later trials (*μ*_Trial 1_ = 0.15 g [90% CI = 0.08 - 0.24]; *μ*_Trial 2_ = 0.17 g [90% CI = 0.10 - 0.28]; *μ*_Trial 3_ = 0.27 g [90% CI = 0.14 - 0.47]). Much of the variation in whether or not an individual woodrat consumed food was explained by whether or not a woodrat had been in multiple trials (σ_order_ = 0.71 [90% CI = 0.02 - 1.47]).

## DISCUSSION

In this study, we investigated the dietary plasticity of dusky-footed woodrats during a time of extreme environmental and ecological stress. Our results demonstrate that during drought conditions, woodrats are consuming high quantities of chamise, a highly chemically-defended, drought-tolerant plant, in addition to their more traditional oak diet (Atsatt and Ingram 1983). Additionally, we found important age-based differences. Adults included more plants in their preferred diet compared to juveniles. Adults preferred four plants in rank order: scrub oak (25.8%), California Buckeye (18.9%), poison oak (17.9%), and chamise (17.6%). In contrast, juvenile preferences were less diverse, including only two plants in rank order: scrub oak (46.2%) and chamise (16.7%). Notably, chamise accounted for similar percentages of the preferred diets of adults and juveniles. The main difference was in how much scrub oak was consumed. Juveniles relied much more heavily on oak compared to adults, who instead ate a wider variety of the available plants. All of these plants are chemically defended with distinct sets of secondary compounds, which can have toxic effects on herbivores and require different sets of detoxification mechanisms (Robbins et al. 1991, Dearing et al. 2000). Juveniles may be less likely to sample larger quantities of other plants because they have not yet developed the physiological mechanisms needed to breakdown a diverse array of secondary compounds (Freeland and Janzen 1974). However, experience with small amounts of plant toxins can induce tolerance (Freeland and Janzen 1974, Fox and Morrow 1981), which potentially explains why adults included more plant variety in their preferred diets. It also may partially explain why individuals who were sampled repeatedly were more likely to sample plant materials in subsequent trials, although habituation to the cafeteria trials and other unmeasured ecological factors cannot be ruled out.

Interestingly, woodrats showed the least amount of preference for incense cedar, the novel food source, and also showed the least amount of difference in amount consumed for adults and juveniles (Fig 1e, Fig 2e, Table 3). While cedar is a natural part of woodrat diets in other parts of the species’ range, this finding is in line with research showing that woodrats tend to avoid novel, chemically defended plants (McEachern et al., 2006). In this case, it is likely that both adults and juveniles at Quail Ridge, who have no previous experience with incense cedar, lack the physiological mechanisms needed to detoxify its cocktail of secondary compounds. In future studies, it would also be interesting to see if woodrats are more likely to sample novel, chemically defended foods in the presence of social information such as olfactory cues, as has been found in Norway Rats (Galef, 1996).

### Edge Effects, Age, and Diet Preferences

Distance from habitat edge and its interaction with woodrat age was one of the most important predictors of diet preference during cafeteria trials. Adults who lived farthest from the habitat edge were more likely to sample both larger quantities and a wider range of plants compared to adults living near the edge. With the exception of scrub oak, adult woodrats furthest from the habitat edge sampled over twice as much of each plant material compared to adults living close to the edge (Figure 3). However, when pooled over all plants, this effect is much less pronounced (Figure 3h). Few adults occupied dens near edge habitat (only 1 adult occupied a den within 10 meters of the edge, compared to 4 juveniles in the same category). Thus the edge effect we observed in adults may be an artifact of sampling variation, or perhaps due to some social or ecological factor we did not measure. More data is needed to clarify whether this result has ecological significance.

Interestingly, juveniles displayed the opposite edge-effect. Juveniles living closest to the edge consumed larger quantities of plants than juveniles living in interior scrub oak habitat (Figure 3). This result may be due to the possibility that juveniles living close to the edge occupy lower quality habitat and are thus more nutritionally stressed than juveniles in the interior. Several factors including increased risk of predation and increased exposure to environmental extremes near edges could contribute to this difference. Another important factor is whether and how far juveniles must disperse from their natal territory. Dispersing juveniles may be more likely to occupy lower quality territories near the edge, and may be more nutritionally stressed due to the energetic costs of dispersal combined with the consequences of residing in lower quality habitat. Other studies have found similar age-based edge effects in small mammals, with adults tending to occupy home ranges in the center of habitat patches and juveniles occupying home ranges near the edge, presumably due to their competitive inferiority (e.g, gray-tailed voles, Lidicker and Peterson 1999).

Both adult and juvenile woodrats showed a strong preference for scrub oak (Fig 1g and Fig 2g). For adults, we found a near zero relationship between distance from den to habitat edge, whereas in juveniles we observed a marked decrease in scrub oak consumption in woodrats occupying dens further from the edge. The overall preference for scrub oak is unsurprising, as it is a common resource in their habitat and is often a highly nutritional and reliable food source. The markedly higher consumption of scrub oak by juveniles closer to the edge might be due to the fact that they are nutritionally stressed and occupy low quality territories.

While this study is not the first to document that woodrat consume chamise (Linsdale and Tevis 1951), we were surprised by the quantities of chamise consumed and its high rank in the preferred diet for adults and juveniles. Our results suggest that woodrats may be actively crossing the edge boundary to access chamise, as it is rare in scrub oak habitat. This result echoes similar observations by Sakai and Noon (1997) in forested habitats. We hypothesized that individuals living closer to edge habitat would be more likely to consume chamise. Our results support this hypothesis, but the effect was largely driven by juvenile food preferences. The opposite edge effect was observed in adult woodrats (Fig 1f and 2f)—adults living *further* from the habitat edge were *more* likely to consume chamise. However, this result is difficult to interpret given that most adults lived in interior scrub oak habitat and not near the edge. For this reason, we cannot rule out the role of sampling variation in driving this result. Additional data would help clarify the interaction between age and distance to edge that we observed.

### Management Implications

Edge habitat presents a tradeoff for wildlife, with both positive and negative effects on populations. There is a rich literature demonstrating the negative effects of habitat edges, which can expose individuals to higher levels of biotic and abiotic stress (Harris 1988, Murcia 1995). However, edge habitat is not universally bad under all contexts, and some species appear to be “edge species” with greater abundance near edges (e.g., deer and taiga voles, Wolf and Lidicker 1980). Among mammals, the cumulative effects of edges appear to be neutral or positive (Kremsater and Bunnell 1999). Clearly, in the context of extreme drought, proximity to natural edge habitat (i.e, chamise chaparral, or chamissal) can be beneficial to woodrat populations at our study site. Woodrat populations living near edges are able to exploit chamise as an alternative food resource and are able to maintain higher density populations compared to habitats lacking access to chamise. Other studies have documented the value of edge habitat to supporting abundant populations of woodrats (Sakai and Noon 1993) and their predators (Sakai and Noon 1997). Our findings suggest that chaparral edge habitats bordering oak woodlands can help sustain woodrat populations, and may provide refugia during extreme or prolonged droughts when preferred resources such as oaks provide a less reliable food source. It is important to consider, however, that other ecological factors may be at play and a broader sampling effort across different sites is necessary to fully examine the robustness of this hypothesis. Other studies have documented declines in woodrat populations related to drought conditions such as reduced rainfall (Gillespie 2008) and aridity (Spevak 1983) that can significantly affect food resources. However, the effects of food limitation are also probably interacting with predation pressures, and territory quality, which have not been quantified at Quail Ridge Reserve. Additional research exploring the tradeoffs between increased access to chamise and increased predation risk is needed to elucidate the importance of edge habitats as refugia for woodrat populations during drought.

An important finding of this study is that woodrats will readily consume large quantities of chamise- a plant that is rare in the woodrat’s preferred scrub oak habitat, and highly chemically defended by plant secondary compounds. This observation is important for several reasons. Nutritional quality and leaf abundance in California oaks is reduced during drought (Callaway & Nadkarni 1991), so it is likely an important survival strategy for woodrats to seek out other food sources when droughts occur. In other mammals, reduced oak nutritional quality during drought has been linked to a reduction in lactation quality and subsequent switch in diet preference (Lashley and Harper 2012). The ranges of many oak species- a preferred food of woodrats across California’s mediterranean ecosystems- are expected to shrink as the climate changes (Kueppers et al. 2005; Loarie et al. 2008) and more drought and fire-tolerant plants, such as chamise may expand their ranges locally.

This study suggests a management plan for woodrats (and for other species that depend upon woodrat dens for shelter) that prioritizes the maintenance of edge habitat, specifically chamissal, in addition to their preferred oak woodland habitat during times of drought. Drought is a fundamental aspect of California’s climate. For the past 5 years and counting, California has been experiencing the worst drought in over a century of observation (Griffin and Anchukaitis 2014). The state’s extreme arid conditions from autumn 2013 through spring 2014, marked some of the lowest water totals in the climate record. Particularly in Central and Southern California, the effects of low precipitation were intensified by record breaking hot temperatures (Griffin and Anchukaitis 2014, Vose et al. 2014, NOAA 2014), and it is only expected to increase the intensity of evaporation and drought in the future (Seager et al. 2007). Due to these current and predicted climatic shifts, it is imperative to monitor and observe ecological responses, in particular that of keystone and indicator species such as woodrats.

## ACKNOWLEDGEMENTS

We thank Honora Tisell, Ken Crews, Colin Turcotte, and Felipe Santich, for field assistance, and Shane Waddel for logistical assistance at Quail Ridge Reserve. In addition, we are thankful to Richard McElreath for providing statistical advice and draft feedback, and Karen Mabry for offering feedback on drafts of this manuscript. Funding was provided by the grants awarded to BJB by the University of California, Davis Natural Reserve System and an ARCS Foundation, Northern California Chapter award. This material is based upon work supported by the National Science Foundation Graduate Research Fellowship under Grant No. 1148897 awarded to BJB. Any opinions, findings, and conclusions or recommendations expressed in this material are those of the authors and do not necessarily reflect the views of the National Science Foundation.

## LITERATURE CITED

Atsatt, P. R. and T. Ingram. 1983. Adaptation to oak and other fibrous, phenolic-rich foliage by a small mammal, Neotoma fuscipes. Oecologia 60:135–142.

Barbour, E. and L. M. Kueppers. 2011. Conservation and management of ecological systems in a changing California. Climatic Change 111:135–163.

Cahill, A. E. et al. 2013. How does climate change cause extinction? Proceedings of the Royal Society of London B: Biological Sciences 280:20121890.

Callaway, R. M. and N. M. Nadkarni. 1991. Seasonal patterns of nutrient deposition in a Quercus douglasii woodland in central California. Plant and Soil 137:209–222.

Caro, T. M. 2010. Conservation by Proxy: Indicator, Umbrella, Keystone, Flagship, and Other Surrogate Species. 2 edition. Island Press, Washington, DC.

Cayan, D. R., E. P. Maurer, M. D. Dettinger, M. Tyree and K. Hayhoe. 2008. Climate change scenarios for the California region. Climatic Change 87:21–42.

Clobert, J., M. Baguette, T. G. Benton, J. M. Bullock and S. Ducatez. 2012. Dispersal Ecology and Evolution. Oxford University Press.

Dearing, M. D., A. M. Mangione and W. H. Karasov. 2000. Diet breadth of mammalian herbivores: nutrient versus detoxification constraints. Oecologia 123:397–405.

Fox, L. a and P. A. Morrow. 1981. Specialization: species property or local phenomenon. Science 211:887–893.

Freeland, W. J. and D. H. Janzen. 1974. Strategies in Herbivory by Mammals: The Role of Plant Secondary Compounds. The American Naturalist 108:269–289.

Fried, J. S., M. S. Torn and E. Mills. 2004. The Impact of Climate Change on Wildfire Severity: A Regional Forecast for Northern California. Climatic Change 64:169–191.

Galef, B. G. 1996. Social Enhancement of Food Preferences in Norway Rats: A Brief Review. in Social Learning In Animals: The Roots of Culture (C. M. Heyes & B. G. Galef, eds.).

Gillespie, S. C., D. H. Van Vuren, D. A. Kelt, J. M. Eadie and D. W. Anderson. 2008. Dynamics of Rodent Populations in Semiarid Habitats in Lassen County, California. Western North American Naturalist 68:76–82.

Giorgi, F. and P. Lionello. 2008. Climate change projections for the Mediterranean region. Global and Planetary Change 63:90–104.

Griffin, D. and K. J. Anchukaitis. 2014. How unusual is the 2012–2014 California drought? Geophysical Research Letters 41:2014GL062433.

Guisan, A. and W. Thuiller. 2005. Predicting species distribution: offering more than simple habitat models. Ecology Letters 8:993–1009.

Harris, L. D. 1988. Edge Effects and Conservation of Biotic Diversity. Conservation Biology 2:330–332.

Howard F. Sakai, B. R. N. 1993. Dusky-footed woodrat abundance in different aged forests in northwestern California. The Journal of Wildlife Management 57.

Kremsater, L. and F. L. Bunnell. 1999. Edge effects: theory, evidence and implications to management of western North American forests. in Forest Fragmentation: Wildlife and Management Implications (J. A. Rochelle, L. A. Lehmann & J. Wisniewski, eds.). BRILL.

Kueppers, L. M., M. A. Snyder, L. C. Sloan, E. S. Zavaleta and B. Fulfrost. 2005. Modeled regional climate change and California endemic oak ranges. Proceedings of the National Academy of Sciences of the United States of America 102:16281–16286.

Lashley, M. A. and C. A. Harper. 2012. The Effects of Extreme Drought on Native Forage Nutritional Quality and White-Tailed Deer Diet Selection. Southeastern Naturalist 11:699–710.

Le Galliard, J.-F., A. Remy, R. A. Ims and X. Lambin. 2012. Patterns and processes of dispersal behaviour in arvicoline rodents. Molecular Ecology 21:505–523.

Leopold, A. 1986. Game Management. Reprint edition. University of Wisconsin Press, Madison, Wis.

Lidicker, W. Z. 1999. Responses of mammals to habitat edges: an overview. Landscape Ecology 14:333–343.

Linsdale, J. M. and L. P. Tevis. 1951. The dusky-footed wood rat: a record of observations made on the Hastings Natural History Reservation. University of California Press.

Loarie, S. R. et al. 2008. Climate Change and the Future of California’s Endemic Flora. PLoS ONE 3:e2502.

Lynch, M. F., A. L. Fesnock and D. V. Vuren. 1994. Home Range and Social Structure of the Dusky-Footed Woodrat (Neotoma fuscipes). Northwestern Naturalist 75:73–75.

Matocq, M. D. 2002. Morphological and Molecular Analysis of a Contact Zone in the Neotoma Fuscipes Species Complex. Journal of Mammalogy 83:866–883.

McEachern, M. B., C. A. Eagles-Smith, C. M. Efferson and D. H. Van Vuren. 2006. Evidence for local specialization in a generalist mammalian herbivore, Neotoma fuscipes. Oikos 113:440–448.

McEachern, M. B., J. M. Eadie and D. H. V. Vuren. 2007. Local genetic structure and relatedness in a solitary mammal, Neotoma fuscipes. Behavioral Ecology and Sociobiology 61:1507–1507.

McEachern, M. B., R. L. McElreath, D. H. Van Vuren and J. M. Eadie. 2009. Another genetically promiscuous “polygynous” mammal: mating system variation in Neotoma fuscipes. Animal Behaviour 77:449–455.

McElreath, R. 2015. rethinking: Statistical Rethinking book package.

Mooney, H. A. and P. W. Rundel. 1979. Nutrient Relations of the Evergreen Shrub, Adenostoma fasciculatum, in the California Chaparral. Botanical Gazette 140:109–113.

Murcia, C. 1995. Edge effects in fragmented forests: implications for conservation. Trends in Ecology & Evolution 10:58–62.

NOAA National Centers for Environmental Information. 2014. State of the Climate: National Overview for August 2014.

R Core Team. 2014. R: A Language and Environment for Statistical Computing. R Foundation for Statistical Computing, Vienna, Austria.

Ries, L. R. J. F. Jr., J. Battin and T. D. Sisk. 2004. Ecological Responses to Habitat Edges: Mechanisms, Models, and Variability Explained. Annual Review of Ecology, Evolution, and Systematics 35:491–522.

Robbins, C. T., A. E. Hagerman, P. J. Austin, C. McArthur and T. A. Hanley. 1991. Variation in Mammalian Physiological Responses to a Condensed Tannin and Its Ecological Implications. Journal of Mammalogy 72:480–486.

Rowe, K. C. et al. 2015. Spatially heterogeneous impact of climate change on small mammals of montane California. Proceedings of the Royal Society of London B: Biological Sciences 282:20141857.

Rundel, P. W. and D. J. Parsons. 1979. Structural Changes in Chamise (Adenostoma fasciculatum) along a Fire-Induced Age Gradient. Journal of Range Management 32:462–466.

Sakai, H. F. and B. R. Noon. 1993. Dusky-Footed Woodrat Abundance in Different-Aged Forests in Northwestern California. The Journal of Wildlife Management 57:373–382.

Sakai, H. F. and B. R. Noon. 1997. Between-Habitat Movement of Dusky-Footed Woodrats and Vulnerability to Predation. The Journal of Wildlife Management 61:343–350.

Seager, R. et al. 2007. Model Projections of an Imminent Transition to a More Arid Climate in Southwestern North America. Science 316:1181–1184.

Sikes, R. S. and W. L. Gannon. 2011. Guidelines of the American Society of Mammalogists for the use of wild mammals in research. Journal of Mammalogy 92:235–253.

Spevak, T. A. 1983. Population Changes in a Mediterranean Scrub Rodent Assembly during Drought. The Southwestern Naturalist 28:47–52.

Stamps, J. A., M. Buechner and V. V. Krishnan. 1987. The Effects of Edge Permeability and Habitat Geometry on Emigration from Patches of Habitat. The American Naturalist 129:533–552.

Stan Development Team. 2015. RStan: the R interface to Stan.

Tuomainen, U. and U. Candolin. 2011. Behavioural responses to human-induced environmental change. Biological Reviews of the Cambridge Philosophical Society 86:640–657.

Varner, J. and M. D. Dearing. 2013. Dietary plasticity in pikas as a strategy for atypical resource landscapes. Journal of Mammalogy 95:72–81.

Vose, R. S. et al. 2014. Improved Historical Temperature and Precipitation Time Series for U.S. Climate Divisions. Journal of Applied Meteorology and Climatology 53:1232–1251.

Watanabe, S. 2010. Asymptotic Equivalence of Bayes Cross Validation and Widely Applicable Information Criterion in Singular Learning Theory. J. Mach. Learn. Res. 11:3571–3594.

Whitford, W. G. and Y. Steinberger. 2010. Pack rats (Neotoma spp.): Keystone ecological engineers? Journal of Arid Environments 74:1450–1455.

Wolff, J. O. and W. Z. Lidicker Jr. 1980. Population ecology of the taiga vole, Microtus xanthognathus, in interior Alaska. Canadian Journal of Zoology 58:1800–1812.

